# Identification of mundulone and mundulone acetate as natural products with tocolytic efficacy in mono- and combination-therapy with current tocolytics

**DOI:** 10.1101/2021.05.13.444040

**Authors:** Shajila Siricilla, Christopher J. Hansen, Jackson H. Rogers, Carolyn L. Simpson, Stacey L. Crockett, Jeff Reese, Bibhash C. Paria, Jennifer L. Herington

## Abstract

Currently, there are a lack of FDA-approved tocolytics for the management of preterm labor. We previously observed that the isoflavones mundulone and mundulone acetate (MA) inhibit intracellular Ca^2+^-regulated myometrial contractility. Here, we further probed the potential of these natural products to be small molecule leads for discovery of novel tocolytics by: (1) examining uterine-selectivity by comparing concentration-response between human primary myometrial cells and a major off-target site, aortic vascular smooth muscle cells (VSMCs), (2) identifying synergistic combinations with current clinical tocolytics to increase efficacy or and reduce off-target side effects, (3) determining cytotoxic effects and (4) investigating the efficacy, potency and tissue-selectivity between myometrial contractility and constriction of fetal ductus arteriosus (DA), a major off-target of current tocolytics. Mundulone displayed significantly greater efficacy (E_max_ = 80.5% vs. 44.5%, p=0.0005) and potency (IC_50_ = 27 μM and 14 μM, p=0.007) compared to MA in the inhibition of intracellular-Ca^2+^ from myometrial cells. MA showed greater uterine-selectivity, compared to mundulone, based on greater differences in the IC_50_ (4.3 vs. 2.3 fold) and E_max_ (70% vs. 0%) between myometrial cells compared to aorta VSMCs. Moreover, MA demonstrated a favorable *in vitro* therapeutic index of 8.8, compared to TI = 0.8 of mundulone, due to its significantly (p<0.0005) smaller effect on the viability of myometrial (hTERT-HM), liver (HepG2) and kidney (RPTEC) cells. However, mundulone exhibited synergism with two current tocolytics (atosiban and nifedipine), while MA only displayed synergistic efficacy with only nifedipine. Of these synergistic combinations, only mundulone + atosiban demonstrated a favorable TI = 10 compared to TI=0.8 for mundulone alone. While only mundulone showed concentration-dependent inhibition of *ex vivo* mouse myometrial contractions, neither mundulone or MA affected mouse fetal DA vasoreactivity. The combination of mundulone and atosiban yielded greater tocolytic efficacy and potency on term pregnant mouse and human myometrial tissue compared to single-drugs. Collectively, these data highlight the difference in uterine-selectivity of Ca^2+^-mobilization, effects on cell viability and tocolytic efficacy between mundulone and MA. These natural products could benefit from medicinal chemistry efforts to study the structural activity relationship for further development into a promising single- and/or combination-tocolytic therapy for management of preterm labor.

**Chemical compounds studied in this article:** atosiban (Pubchem CID: 5311010); indomethacin (Pubchem CID: 3715); mundulone (Pubchem CID: 4587968); mundulone acetate (Pubchem CID: 6857790); nifedipine (Pubchem CID: 4485); oxytocin acetate (Pubchem CID: 5771); U46619 (Pubchem CID: 5311493)

## 1. Introduction

Preterm birth (PTB) rates continue to rise with over 15 million cases/year globally and remains the greatest contributor to neonatal morbidities and mortalities [1, 2]. The causes of PTB are multifactorial, yet despite the initial trigger, stimulation of intracellular Ca^2+^-release in myometrial cells is the final common pathway controlling uterine contractions [3, 4]. Current clinically-utilized drugs to manage preterm labor include: nifedipine (calcium channel blocker), indomethacin (cyclooxygenase inhibitor) or terbutaline (β-adrenergic agonist). However, these drugs are not approved by the FDA for tocolytic use due to their detrimental off-target side effects for either infant or mother, and short duration of benefit. Specifically, these drugs are known to cause either maternal cardiovascular effects, *in utero* constriction of the fetal ductus arteriosus (DA) or fetal tachycardia [5-7]. Atosiban (oxytocin receptor antagonist) is a safe tocolytic without serious side-effects but has not been granted FDA-approval due a lack of clinical efficacy [8, 9]. Thus, novel safe and effective tocolytic agents are urgently needed for management of preterm labor. Furthermore, due to the multiple pathways involved upstream of intracellular-Ca^2+^ to regulate myometrial contractility, combining current off-label tocolytics with small-molecules targeting different molecular signaling pathways may provide a synergistic effect, permitting lower doses that reduce off-target effects or increase therapeutic efficacy.

In our continued efforts to discover novel tocolytics, we have previously performed a pilot screen of small molecules from the Spectrum collection of compounds comprised of drug components, natural products and other bioactive molecules with a wide range of biological activity [3]. A high-throughput assay was used to screen small-molecules and identified the isoflavone natural-product mundulone and its derivative mundulone acetate (MA) (Fig.1) as potent hit-antagonists of Ca^2+^-mobilization from primary myometrial cells. Mundulone is extracted from the bark of *Mundulea sericea* and its structure was elucidated in 1959 to be of the isoflavone subclass of flavonoids [10]. Isoflavones have several pharmacological effects, including smooth muscle contractions in the uterus and vasculature [11-15]. Moreover, a recent review on natural products for tocolysis found that most plant extracts for inhibition of uterine contractions belong to flavonoid or terpene classes [16]. Interestingly, plant extracts containing flavonoids have shown tocolytic effects (*J. flava, B. pinnatum, T. paniculatum*), with *B. pinnatum* entering into a clinical trial that was, unfortunately, withdrawn early due to lack of patient enrollment [17-22]. Therefore, mundulone and MA belong to an interesting class of compounds to study for tocolytic potential.

**Figure 1.**
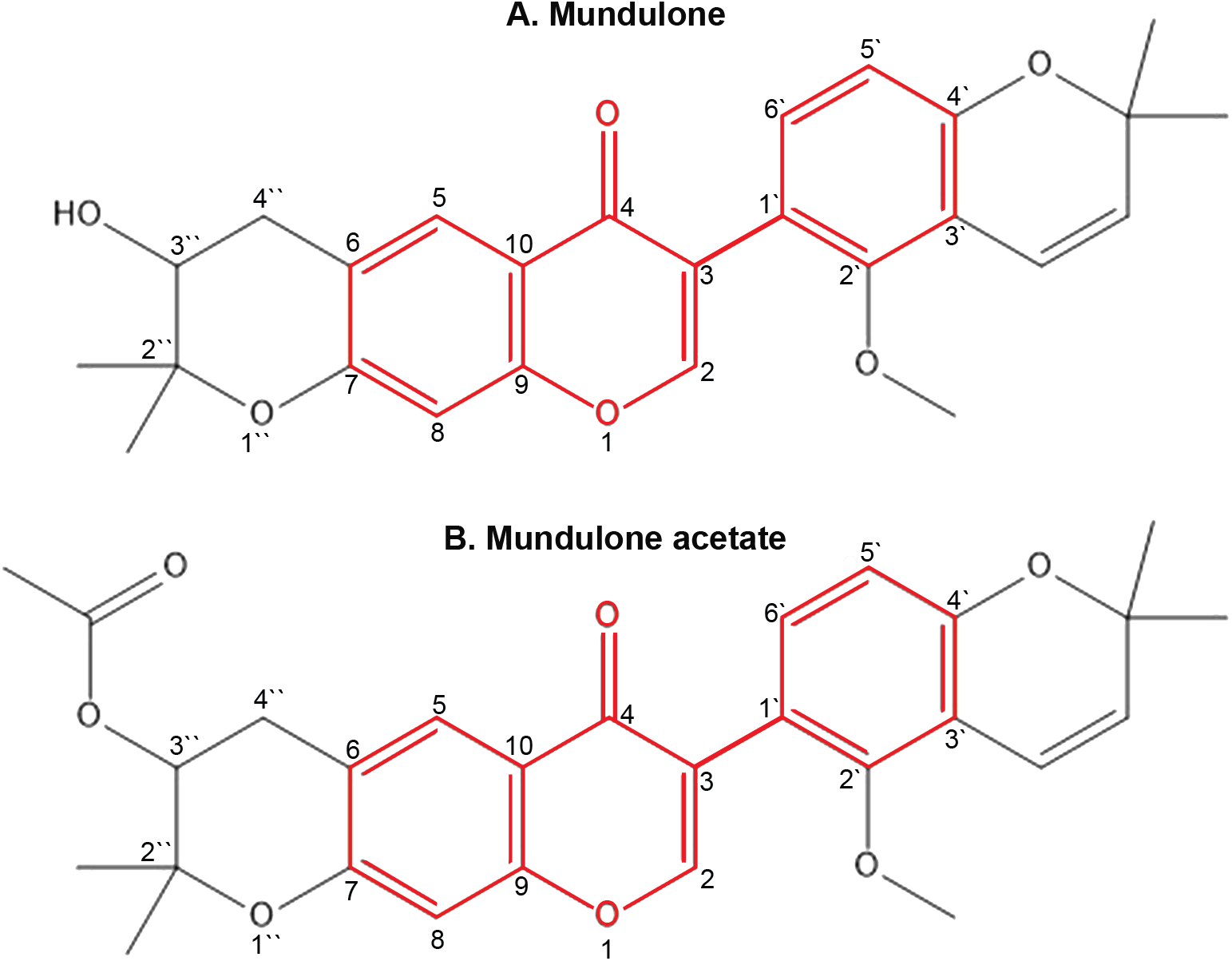
Structures of mundulone and MA. The isoflavone group of each compound is highlighted.

Based on our prior discovery that mundulone and MA to inhibit *in vitro* myometrial Ca^2+^-mobilization and the reported tocolytic ability of other isoflavones, we aimed to further probe mundulone and MA for tocolytic drug development. The goal of this work was to: 1) examine the uterine-selectivity Ca^2+^-mobilization inhibition by comparative testing vascular smooth muscle cells (VSMCs) - the major off-target limiting the use of current tocolytics, 2) determine cytotoxic effects on myometrial cells and metabolic organ (kidney and liver) cells, 3) identify synergistic combinations of mundulone and MA with current tocolytics to increase efficacy and/or potency for reduced off-target side effects and 4) determine the *ex vivo* tocolytic efficacy and potency on mouse myometrial tissue and confirm uterine selectivity at the tissue level by evaluating their effect on the *ex vivo* constriction of fetal ductus arteriosus (DA), another major off-target of known tocolytics.

## 2. Materials and methods

### 2.1. Compounds and drugs

Drugs and compounds were procured from the following vendors: MicroSource Discovery Systems [mundulone (00200011) and MA (00200019)], Sigma [atosiban (A3480) and oxytocin acetate (O6379)] and Cayman Chemicals [nifedipine (11106), indomethacin (70270) and U46619 (16450].

### 2.2. Primary human uterine myometrial tissue and cells

Human myometrial tissue biopsies were obtained at the time of cesarean section under the Vanderbilt University Institutional Review Board protocol #150791. Reproductive-age (18-45 years) women undergoing a scheduled or repeat cesarean section at term gestation (≤ 39 weeks) were recruited, fully informed and consented to the study. Inclusion criteria comprised of breech presentation, previous cesarean section, fetal anomaly, fetal distress or placental previa. Exclusion criteria included: clinical or histological signs of vaginal/chorioamniotic/intrauterine inflammation/infection, or current use of vasopressors or bronchodilators. Samples were placed into either sterile cold 1XHBSS or 1XPBS and transported to the laboratory for immediate myometrial cell isolation or use in *ex vivo* organ bath studies, respectively, after removal of perimetrium and decidua.

Primary myometrial cells were isolated, selectively enriched for smooth muscle cells and cultured as previously described [3, 23]. Primary myometrial cells were unpassaged and became near-confluent 10-15 days post-isolation, with media (DMEM supplemented with 10% fetal bovine serum (FBS), 25 mM HEPES, 100 U/ml penicillin-streptomycin) changed every 3 days. The cells were dissociated at near-confluency using 0.25% Trypsin-EDTA, and then plated at 4,000 cells/well in black-walled 384-well plates (Grenier Bio-One) for Ca^2+^-mobilization assays exactly as previously described [23].

### 2.3 Aortic vascular smooth muscle cells (VSMCs)

Human primary aortic VSMC were purchased from ATCC (ATCC® PCS-100-012™), cultured in complete vascular cell growth medium (vascular cell basal medium (ATCC® PCS-100-030™ supplemented with 5 ng/mL rh FGF-basic, 5 µg/mL rh Insulin, 50 µg/mL ascorbic acid, 10 mM L-glutamine, 5 ng/mL rh EGF, 5% fetal bovine serum and 100 U/mL penicillin-streptomycin) and passaged at ∼80% confluency. Aortic VSMCs from passage 3 were plated at 4,000 cells/well in similar 384-well plates described above for Ca^2+^-mobilization assays.

### 2.4. Comparative concentration-dependent Ca^2+^-mobilization

Mundulone and MA were assayed at 10-point three-fold titrations starting at 60μM in triplicate, as previously described [3], to compare the E_max_ and IC_50_ values between myometrial and VSMCs. A dual-addition Ca^2+^-mobilization assay was performed exactly as previously described using the automated equipment within the Vanderbilt University High-Throughput Screening Facility [3, 23]. Briefly, Fluo-4AM (2μM) was loaded into cells and a Panoptic (WaveFront Biosciences) integrated pipettor was used to add the titrations of mundulone or MA to cells. After 30 minutes of incubation, baseline fluorescence was measured for 20sec, followed by the addition of the EC_80_ value of either oxytocin (OT) or U46619, and the fluorescence (Ca^2+^-mobilization) was continuously measured in real time for an additional 120sec. Oxytocin was utilized as an agonist during the second addition for myometrial cells, while U46619 (a thromboxane A_2_ receptor agonist) was used for aorta VSMCs. On each day the assay was performed, the EC_80_ values of OT and U46619 were calculated from 13-point, three-fold titrations (starting at 100μM) to determine the ability of mundulone and MA to inhibit this submaximal concentration. All concentration response curves (CRCs) were performed in triplicate. For each well, the mean baseline value (MBV) for Ca^2+^-fluorescence was calculated during the 0–19 sec timeframe. The max relative fluorescent unit (RFU) was calculated from the 20–140 sec timeframe, and the baseline was subtracted from the max RFU. An average Max-MBV RFU was calculated for each concentration of agonist (OT and U46619). Data were analyzed using WaveGuide software and the % inhibition for each concentration of mundulone and MA was calculated. Non-linear regression analyses using Graphpad Prism 8.0 were performed to generate CRCs and to determine the E_max_ and IC_50_ values. Fold difference in efficacy was calculated as E_max_ (maximum % inhibition achieved by the compound) on myometrial cells plus the absolute E_max_ on aorta VSMCs (E_max-myo_ - E_max-aorta VSMC_). Conversely, the fold difference in potency was calculated as the ratio of IC_50_ (concentration of the compound required to achieve 50% of Emax) of each compound on aorta VSMCs to the IC_50_ on myometrial cells (IC_50-aorta_/IC_50-myo_). Two-way analysis of variance followed by a post hoc Fisher’s LSD test was used to determine significant differences between the E_max_ values.

### 2.5. Combination high-throughput Ca^2+^-mobilization assay

An 8×8 dose matrix was used to evaluate the combination effects between two serially-diluted single-compound concentrations. Mundulone and MA were combined with known tocolytics: atosiban, indomethacin and nifedipine. The single-compound IC_50_ values were utilized with three 2-fold titrations above and below the IC_50_ value. Control (no compound) additions were included in the dose matrix. Moreover, the first row and the first column of each matrix contained the individual compound concentrations used to compare the effect of combination. The high-throughput Ca^2+^-assay was performed in 384-well format, allowing for 6 combinations per plate, using the methods described above. Precise titrations of each compound into a compound plate were performed using an Echo555 within the Vanderbilt High-Throughput Screening Facility. To account for position bias and edge effects, the matrices were repeated at different positions of the 384-well plate. Also, the z’-factor determined during the optimization of the Ca^2+^ assay [3, 23] ensures lack of position effects. Raw data was first analyzed using Waveguide, as described above and previously reported [23]. After calculating the %Response for each compound concentration examined, we determined synergistic combinations using Combenefit software [24], which provided synergy scores and heat-maps to visualize models of Bliss independence, Highest Single Agent and Loewe additivity. After selecting up to 3 fixed ratios of synergistic drug combinations, a concentration-response analysis was performed as described in detail above, to confirm whether the synergy was a result of increased efficacy and/or potency.

### 2.6. Cell viability assay

Cytotoxicity of single- and combination-compounds on myometrial (hTERT-immortalized human myometrial, hTERT-HM), kidney (human primary renal proximal tubule epithelial cells, RPTEC) and liver (human hepatocellular carcinoma, HepG2) cells were assessed using a standard WST-1 (Roche) cell viability assay. hTERT-HM (kindly provided by Dr. Jennifer Condon, Wayne State University), RPTEC (ATCC® PCS-400-010™) and HepG2 (ATCC® HB-8065™) cells were cultured in complete DMEM/F-12 (Gibco 11-039-02), RPTEC media (ATCC® PCS-400-030™) and EMEM (ATCC® 30-2003™), respectively. DMEM/F-12 and HEPG2 media were supplemented with 10% FBS and 1% penicillin-streptomycin, while RPTEC media was supplemented with triiodothyronine, rh EGF, hydrocortisone hemisuccinate, rh Insulin, epinephrine, L-Alanyl-L-Glutamine and transferrin (ATCC® PCS-400-040™). Two-fold serial dilutions of each single-compound were tested at concentrations ranging from 0.78–200 μM (a final volume of 100uL). Additionally, compound combinations at their synergistic fixed ratio (FR) and matched single-compound concentrations were tested. A vehicle (DMSO) and positive control (napabucasin, a compound with known toxicity) was included on each plate. Cells in 96-well cell culture plates (4 × 10^4^ cells per well) using the respective media were incubated with compounds for 72hrs, after which 10uL of WST-1 was added to each well. Absorbance at 450nm and 600nm was read 2hrs after incubation with WST-1. After deduction of background absorbance, the % inhibition in cell viability was calculated. Graphpad Prism 8.0 was used to perform non-linear regression analysis for generation of CRCs for each single-drug and drug combination, as well as determine IC_50_ values. An *in vitro* therapeutic index (TI) was calculated as a ratio of the smallest IC_50_ value among the three cell types (concentration of the compound required to affect 50% of cell viability) to the IC_50_ value from Ca^2+^ mobilization assay [25, 26].

### 2.7. Mouse tissue sample collection

Animal experiments were approved by the Vanderbilt University Institutional Animal Care and Use Committee and conformed to the guidelines established by the National Research Council Guide for the Care and Use of Laboratory Animals. Adult (8–12wk) CD-1® IGS (Charles River Laboratories) mice were housed in a 12h light: 12h dark cycle, with free access to food and water. Timed overnight breedings were performed, and the presence of a copulatory plug was considered day 1 of pregnancy, with the expected date of delivery on the evening of day 19. Mice were euthanized with an overdose of isoflurane, followed by cervical dislocation. The uterus was excised on the morning of day 19 of pregnancy to obtain myometrial strips for *ex vivo* contractility studies, as well as the collection of fetal DA tissue for *ex vivo* myography experiments described below.

### 2.8. *Ex vivo* myometrial contractility assay

An isometric contractility organ bath assay was performed as previously described [3]. Briefly, mouse and human myometrial strips (12mm X 5mm X 1 mm) were attached via silk thread to stainless steel hooks connected to a Radnoti LLC force transducer at one end, while the other end of the tissue was anchored to a glass rod at the base of the tissue bath. Preparations were submerged in a heated and oxygenated (37°C, 95% O2-5% CO2) Radnoti LLC tissue bath containing Kreb’s Bicarbonate Solution (in mM: 136.7 NaCl, 4.7 KCL, 2.5 CaCl_2_2H2O, 1.5 MgCl_2_6H2O, 1.8 NaH_2_PO_4_H2O, 15 C_6_H_12_O_6_ and 2.52 NaHCO_3_). Each mouse strip was placed under 1g tension and allowed to equilibrate in the organ bath for 60min prior to recording baseline spontaneous contractile activity. Only myometrial strips that produced rhythmic contractions were used for the study.

Stock 0.1M Mundulone and Mundulone Acetate was dissolved in 100% PEG-400, while 0.027M of Atosiban was also dissolved in PEG-400. Following the establishment of rhythmic spontaneous contractions, cumulative concentrations (1nM to 0.1mM) of mundulone, MA or vehicle control (PEG-400) were added to individual organ baths every 10 min. In a second set of experiments, cumulative concentrations (1nM to 0.1mM) of a mundulone + atosiban combination at a FR 3.7:1, as well as mundulone and atosiban alone (at their respective concentrations reflective of the ratio) were added to individual organ baths every 10 min or 20 mins depending on whether mouse or human tissue was utilized for the experiment, respectively. After the highest concentration of drug or vehicle examined, tissue viability was determined via exposure to 75mM KCl.

Isometric contractions were recorded using PowerLab/8 SP (ADInstruments) equipment and analyzed with LabChart 7 Pro software (ADInstruments). Contractile activity was assessed by AUC/duration (area under the curve, which is the sum of the integrals for each contraction divided by the ∼600sec duration of each treatment period assessed). All treatment data were then expressed as a percentage of the baseline spontaneous contractile activity. Data were analyzed using Graphpad Prism software and are expressed as mean±SEM (N ≥ 8 uterine strips from 6-9 different mice). Non-linear regression analyses were performed to generate CRCs for calculation of IC_50_ and E_max_. Comparisons of fit were performed to determine if the three-parameter non-linear log fit lines were significantly different (p<0.05) between: 1) mundulone, MA and the vehicle control, as well as 2) mundulone alone, atosiban alone and the mundulone + atosiban combination. Two-way analysis of variance followed by a *post hoc* Tukey test for multiple comparisons was used to determine significant differences between the % response for each concentration of compound.

### 2.9. *Ex vivo* fetal DA myography

The ductus vessels from 7–9 fetal mice, representing at least three different litters, were used for each myography study. The vasoreactivity was evaluated using cannulated, pressurized vessel myography and computer-assisted videomicroscopy, as previously described [31, 32]. Briefly, the ductus was mounted in custom myography chambers (University of Vermont), then equilibrated for 40 mins in modified deoxygenated Krebs buffer at 37°C under 5mmHg of distended pressure. Myography chambers were placed on an inverted microscope equipped with a digital image capture system (IonOptix; Milton, MA) to record changes in the intraluminal vessel diameter. In order to determine vessel viability and peak contractility, the pressure was increased in 5-mmHg increments to 20mmHg, followed by treatment with 50mM KCl deoxygenated modified Krebs buffer (in mM: 64 NaCl, 50 KCl, 2.5 CaCl_2_2H_2_O, 0.9 MgSO_4_, 1 KH_2_PO_4_, 11.1 C_6_H_12_O_6_, 34 NaHCO_3_ (pH 7.3). The vessels were then changed from a flow-through system to a recirculating system (20mL total volume) and allowed to equilibrate for an additional 20 minutes. Baseline lumen diameter was recorded prior to adding cumulative concentrations (1nM to 0.1mM) of mundulone, MA, or vehicle control (DMSO) every 20 mins. After the highest concentration of drug or vehicle examined, vessel viability was determined via exposure to 50mM KCl. All treatment data were expressed as a percent change in baseline lumen diameter. Data are expressed as mean±SEM and non-linear regression analyses in Graphpad Prism 8.0 were performed to generate CRCs for calculation of IC_50_ and E_max_. Comparisons of fit were performed to determine if the non-linear fit lines between mundulone or MA were significantly different (p<0.05) from vehicle control. Two-way analysis of variance followed by a *post hoc* Tukey test for multiple comparisons was used to determine significant differences between the % response for each concentration of compound.

## 3. Results

### 3.1. Uterine-selectivity of mundulone and mundulone acetate

As previously stated, off-target side effects of current off-label tocolytics on maternal cardiovascular and the fetal DA have been major limitations of their use in the management of preterm labor. In a previous study, we discovered that the transcriptome profile of primary human aorta VSMCs was more similar to that of human primary myometrial cells, compared to fetal DA cells, thus serving as useful cells in assays to determine compound uterine-selectivity [23]. To this end, we used our previously established Ca^2+^-mobilization assay to examine whether mundulone and MA were uterine-selective based on either their lack of activity in VSMCs or ≥ 5-fold effect on Ca^2+^-mobilization in myometrial SMCs compared to VSMCs. As shown in Fig.2, mundulone was able to inhibit Ca^2+^-mobilization from myometrial cells and aorta VSMCs, while MA only inhibited Ca^2+^-mobilization in myometrial SMCs (Fig.2 A, C). Thus, MA displayed a significant (p=0.03) 70% difference in efficacy (E_max_), but an insignificant (p=0.37) 4.3-fold difference in potency (IC_50_), between myometrial cells compared to aorta cells (Fig.2D and Table 1). Conversely, mundulone exhibited a significant (p<0.0001) 2.3-fold difference in potency, with no significant (p=0.91) difference in E_max_ (Fig.2B). While both mundulone and MA displayed uterine-selectivity in their potency and efficacy, respectively, there was a significant difference between their efficacy (E_max_ = 80.5% vs. 44.5%, p=0.0005) and potency (IC_50_ = 27 μM and 14 μM, p=0.007) to inhibit Ca^2+^-mobilization in myometrial cells.

**Table 1:** Uterine-selectivity of mundulone and MA.

|  | IC50, uM |  |  |  | Emax, % |  |  | p value |
| --- | --- | --- | --- | --- | --- | --- | --- | --- |
|  | Myometrial | Aorta | Fold difference | p value | Myometrial | Aorta | % difference |  |
| Mundulone | 26.59 | >60 | >2.3 | <0.0001 | 80.52 | 80.15 | 0.37 | 0.91 |
| MA | 13.91 | >60 | >4.3 | 0.37 | 44.45 | -25.61 | 70.06 | 0.03 |

**Figure 2.**
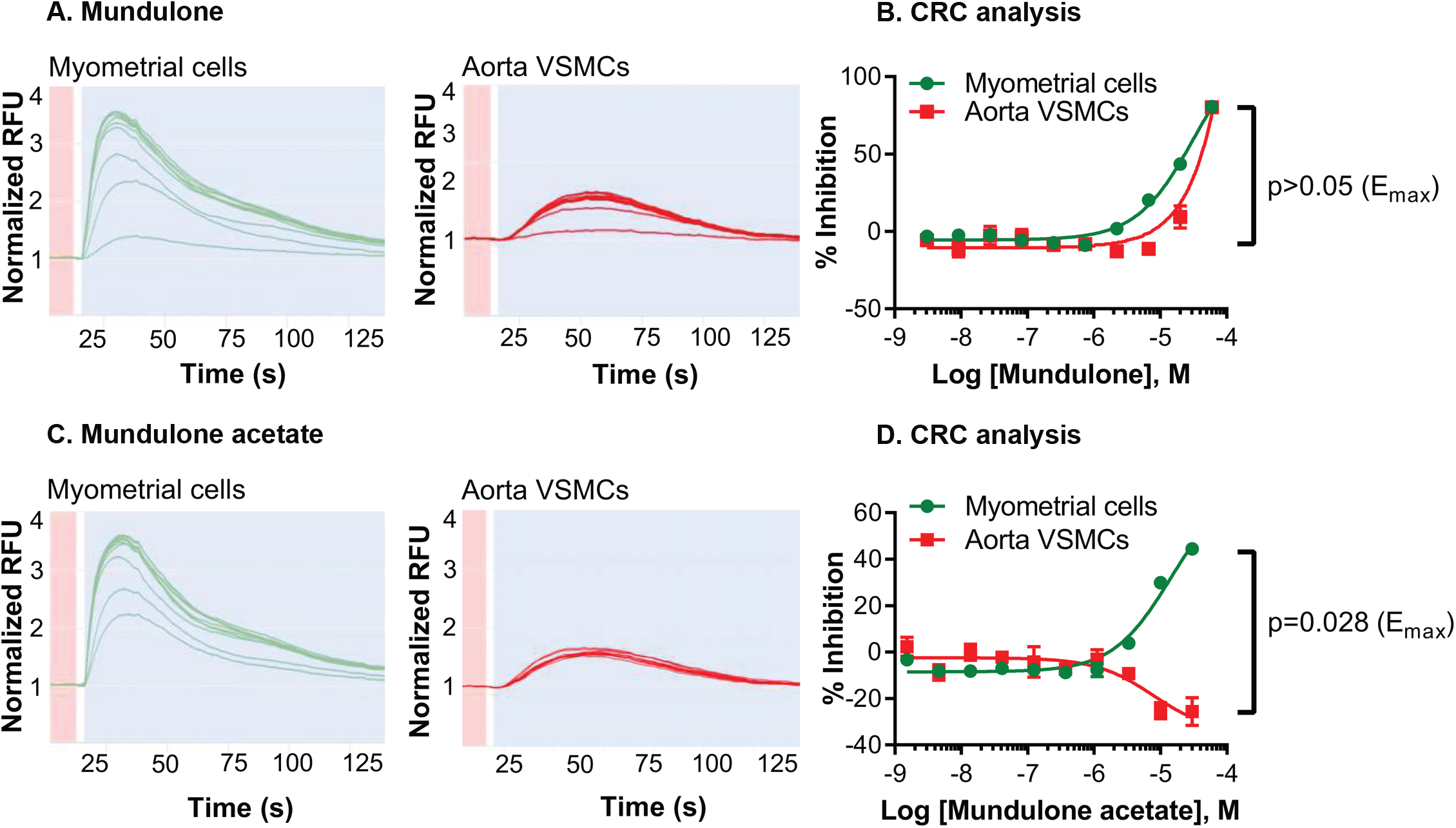
*In vitro* uterine-selectivity of mundulone and MA to inhibit intracellular Ca^2+^-release. Real-time recording of concentration-dependent inhibition of intracellular Ca^2+^-release by mundulone (A) or mundulone acetate (C) from myometrial cells and aorta VSMCs. Concentration-response curves of mundulone (B) or mundulone acetate (D) showing selectivity towards myometrial cells compared to the aorta VSMCs due to either a significant shift in potency (IC_50_) or efficacy (E_max_), respectively. Non-linear regression was used to fit the data (Mean ± SEM) and to calculate the IC_50_ and E_max_., which are provided in Table 1, along with p-values. A 2-way ANOVA with a post-hoc Fisher’s LSD test was used to compare the E_max_ values (shown).

### 3.2. *Ex vivo* tocolytic ability and lack of off-target effect on fetal ductus arteriosus

Based on the *in vitro* inhibition of Ca^2+^-mobilization from the myometrial cells and *in vitro* uterine-selectivity compared to aortic VSMCs by mundulone and MA, we chose to examine their tocolytic ability and confirm uterine-selectivity at the tissue level. A well-established and widely utilized *ex vivo* isometric contractility assay was used to examine the effect of mundulone and MA on contractile activity of term pregnant mouse myometrial tissue [3, 33-39]. During our initial experiments, we observed the precipitation of mundulone and MA from a DMSO stock solution at higher concentrations tested in an aqueous organ bath assay (data not shown). We performed an extensive study to improve the solubility of these, and other compounds, by examining other formulation methods (e.g. alternate solvents, cosolvents, surfactants, complexion and emulsification) [30]. We found that mundulone and MA in an emulsion permitted stable solubility in the organ bath for the duration of the assay. Moreover, the emulsion base was not found to alter the tocolytic efficacy and potency of nifedipine as its formulation in solvent alone [30].

Spontaneous mouse myometrial contractions were recorded (measured in grams, g, of tension), followed by the addition of cumulative concentrations of compounds and vehicle control, as shown in Fig.3 A-B. Mundulone, but not MA, displayed concentration-dependent inhibition of uterine contractions (Fig.3B). The tocolytic efficacy for mundulone (E_max_ = 68%, p=0.009), but not MA (E_max_ = 12%, p=0.43), was significantly greater than the vehicle control (E_max_ = 2%). The potency of mundulone to inhibit *ex vivo* mouse myometrial contractions was 13 uM compared to >0.1mM of MA.

**Figure 3.**
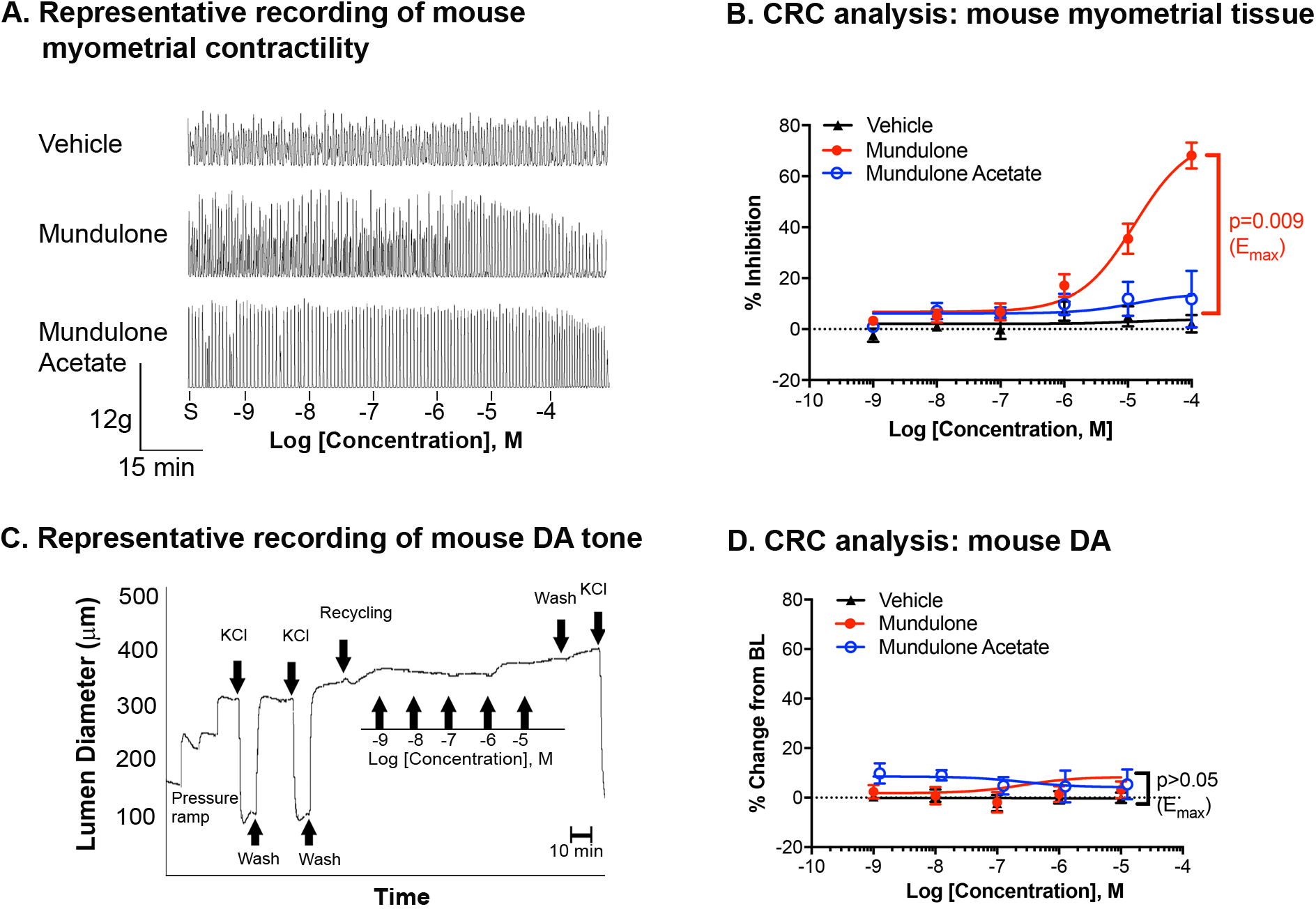
*Ex vivo* tocolytic ability and uterine-selectivity. A. Representative recording of isometric spontaneous contractility (measured in grams of tension) of mouse myometrial tissue prior to the addition of increasing concentrations (10pm - 100uM) of vehicle (emulsion base) control, mundulone or mundulone acetate. B. Recordings were analyzed for contractile AUC. Non-linear regression was used to fit the data (Mean ± SEM) and to calculate IC_50_ and E_max_. C. Representative tracing of fetal mouse ductus arteriosus (DA) tone (measured by lumen diameter) prior to and after the addition of increasing concentrations (1nm - 10uM) of vehicle control, mundulone or mundulone acetate. D. Recordings were analyzed for % change from baseline lumen diameter. A 2-way ANOVA with a post-hoc Tukey analysis was used to compare the E_max_ values (shown).

In order to confirm uterine selectivity at a tissue level, we tested mundulone and MA for their effect on mouse fetal DA vasoreactivity. Neither mundulone or MA affected the vessel diameter of the DA lumen beyond that of the vehicle control (∼10% difference from baseline; Fig.3 C-D, p>0.05). Therefore, selectivity towards uterine myometrial tissue was observed with mundulone and MA inhibiting uterine contractility without causing fetal DA constriction.

### 3.3. *In vitro* synergism with current tocolytics

In order to improve the efficacy and/or potency of mundulone and MA, we explored the possibility of synergism when combined with current tocolytics, which either lack FDA-approval for tocolytic use due to poor side-effects (nifedipine, indomethacin) or efficacy (atosiban). It’s important to note that while these drugs function through different molecular targets, the exact mechanism(s) of action of mundulone and MA are currently unknown. We tested the synergistic potential in our *in vitro* high-throughput Ca^2+^-mobilization assay, which was adapted for compound combination testing using 8×8 dose matrices (Fig.4A). We found that mundulone displayed synergism with atosiban and nifedipine (Fig.4 B-C), while MA exhibited synergistic efficacy with only nifedipine (Fig.4D). Synergistic analysis using the Combenefit online tool for BLISS independence, HSA and Loewe’s additivity models revealed that combination matrices for mundulone + atosiban and mundulone + nifedipine showed several concentration ratios of compounds a and b resulting in synergistic efficacy and potency of either compound a or b. However, the combination matrix for MA and nifedipine displayed only synergistic efficacy. No synergy was observed in mundulone + indomethacin, MA + indomethacin and MA + atosiban combinations (Suppl Fig.1).

**Figure 4.**
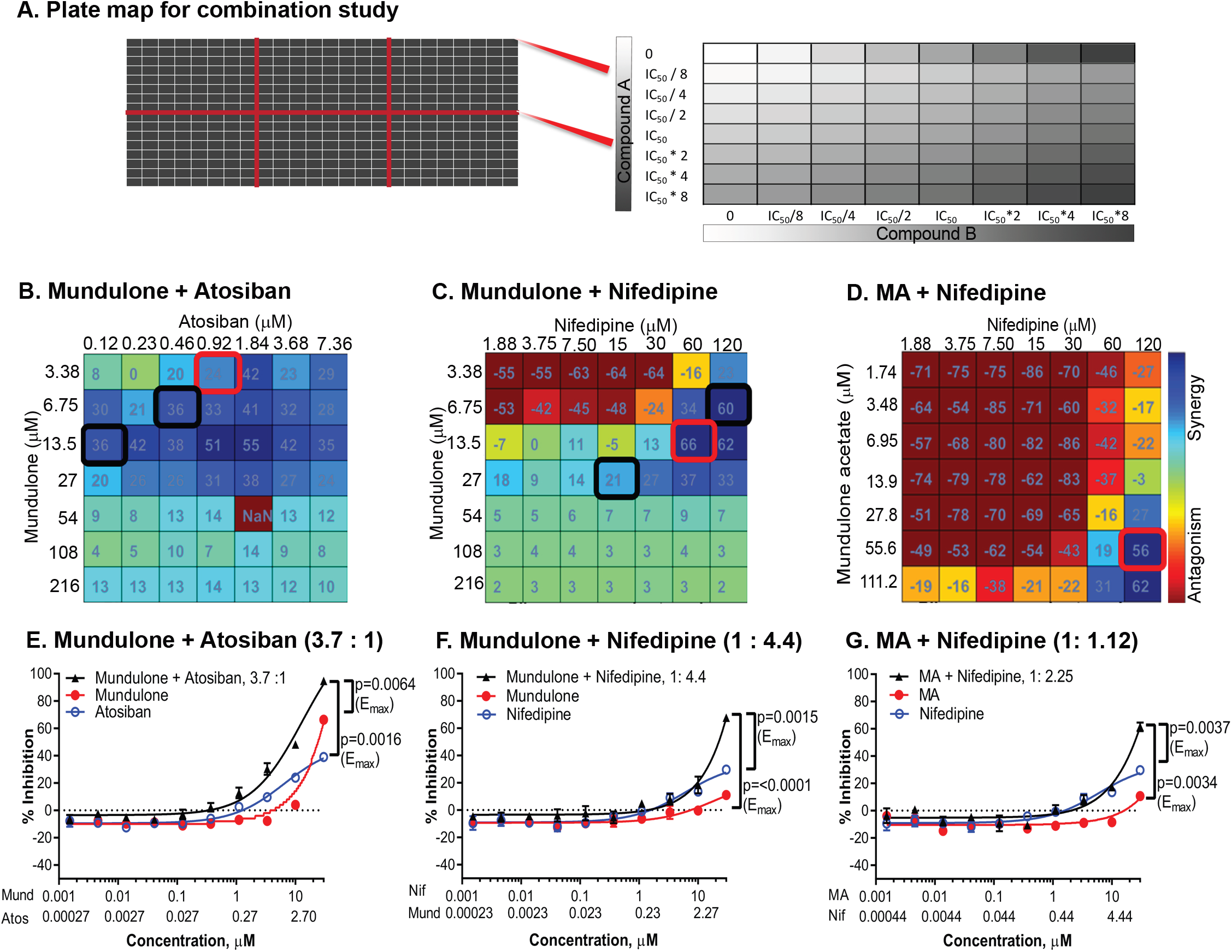
Identification of mundulone and MA synergistic combinations with clinical tocolytics. A. Plate map used for the high-throughput combination Ca^2+^-assay in which one 384-well plate allows testing of six different combinations of two-compounds. Controls (no compounds, white squares) and individual compounds A (columns 1, 9 and 17) and B on (rows H and I, respectively) were included on each plate. The direction of the gradient shows the increasing concentrations of compound A and B. The %response data for each compound concentration was averaged and then analyzed with Combenefit software to provide synergy scores using the Bliss-independence model. Heat maps of synergy scores are shown (B-D). Red or black boxes indicate the three fixed ratios chosen for CRC analysis to confirmation synergistic potency or efficacy. Concentration-response curves of combinations at fixed ratio indicated in red box is shown (E-G). A 2-way ANOVA with a post-hoc Tukey analysis was used to compare the E_max_ values (shown).

For confirmation of synergistic efficacy and/or potency, we performed concentration-response analysis of fixed-ratios of selected combinations (Fig.4 E-G). We picked at least three fixed ratios of drug A and B from the matrix obtained from the high-throughput combination screen. The combination of mundulone + atosiban resulted in both increased efficacy and potency and for all tested ratios, while only increased efficacy was confirmed for mundulone + nifedipine and MA + nifedipine synergistic combinations. The E_max_ and IC_50_ obtained from CRCs of individual compounds in comparison with FRs of compounds in synergistic combinations are listed in Table 2, as are the results of their statistical comparisons. The potency of mundulone improved by a similar degree (2.3 vs 2.7 fold) when in combination with either atosiban (FR 3.7:1) or nifedipine (FR 1:4.4) compared to mundulone as a single-agent. However, the efficacy of mundulone was more greatly improved when in combination with nifedipine than atosiban (6-fold versus 2-fold, respectively), compared to mundulone as a single-agent. Mundulone and mundulone acetate exhibited similar degrees of enhancement in potency (2.7 vs 2.3 fold) and efficacy (6.1 vs 5.8 fold) when in combination with nifedipine.

**Table 2:** Synergistic efficacy and /or potency of fixed-ratio combinations of mundulone and MA with current tocolytics.

| Compound A + Compound B | Ca<sup>2+</sup> assay, IC<sub>50</sub> (μM) |  |  | p value |  | Ca<sup>2+</sup> assay, E<sub>max</sub> (% inhibition) |  |  | p value |  |
| --- | --- | --- | --- | --- | --- | --- | --- | --- | --- | --- |
|  | Compound A | Compound B | Combination*<br>(Compound A;<br>Compound B) | Combination vs<br>Compound A | Combination vs<br>Compound B | Compound A | Compound B | Combination | Combination vs<br>Compound A | Combination vs<br>Compound B |
| Mundulone + Atosiban 112.5 : 1 | >30 | 7.4 | **>30; >0.27** | >0.05 | >0.05 | 66.3 | 4.55 | **93.66** | 0.0082 | <0.0001 |
| Mundulone + Atosiban 14.7 : 1 | >30 | 12.99 | **25.95; 1.77** | >0.05 | >0.05 | 66.3 | 22.59 | **94.62** | 0.0016 | <0.0001 |
| Mundulone + Atosiban 3.7 : 1 | >30 | 6.44 | **13.25; 3.58** | <0.0001 | 0.064 | 66.3 | 39.08 | **94.72** | 0.0064 | 0.0016 |
| Mundulone + Nifedipine 1 : 4.4 | 18.22 | 6.02 | >6.82; >30 | >0.05 | 0.0004 | 11.08 | 29.74 | **67.63** | <0.0001 | 0.0015 |
| MA + Nifedipine 1 : 2.25 | >30 | 6.02 | >13.33; >30 | >0.05 | 0.0015 | 10.68 | 29.74 | **61.47** | 0.0034 | 0.0037 |

Bolded values represent synergistic combinations in potency (IC<sub>50</sub>) and/or efficacy (E<sub>max</sub>). \*Compound A + Compound B values represent the concentration of compound A and compound B present in the combination at 50% inhibition of Ca<sup>2+</sup> mobilization

### 3.4. Cytotoxic effects on myometrial cells and metabolic organ cells

Prediction of *in vivo* cellular toxicity through *in vitro* cell viability assays using a suitable cell type is a key component in early drug discovery. In order to detect the likelihood of cellular toxicity in the target tissue (uterine myometrium) and important xenobiotic metabolic organs (liver and kidney), we assessed the toxic effect of individual compounds mundulone and MA, as well as their synergistic combinations with current tocolytics, through a well-established WST-1 cell viability assay. We used a well characterized immortalized human myometrial (hTERT-HM) cell line for the effects of toxicity on human uterus [23, 40, 41]. HepG2 cells and RPTECs are regularly used as a surrogate for effects of toxicity on human liver and kidney [42-47].

Mundulone had a significantly greater (p<0.0005) effect on the viability of hTERT-HM, HepG2 and RPTEC cells compared to MA (Suppl. Fig.2 and Suppl. Table 1). In relation to the potency of mundulone and MA in the Ca^2+^-assay, MA demonstrated a favorable *in vitro* TI =8.8, while the TI of mundulone was only 0.8. Suppl Table 1 lists the IC_50_ and TI for mundulone and MA for each cell type examined.

The fixed ratios of synergistic combinations tested above were assessed for their effect on cell viability (Fig.5 A-C). The only synergistic combination demonstrating a favorable TI = 10 was that of mundulone + atosiban, which was a great improvement from the TI=0.8 of mundulone as a single agent. Unfortunately, the TI of MA in combination with nifedipine was much lower compared to MA as a single agent (1.8 vs 8.8, respectively). Table 3 lists the IC_50_ and TI for all compound combinations examined, as well as the IC_50_ for their respective single-drug controls.

**Figure 5.**
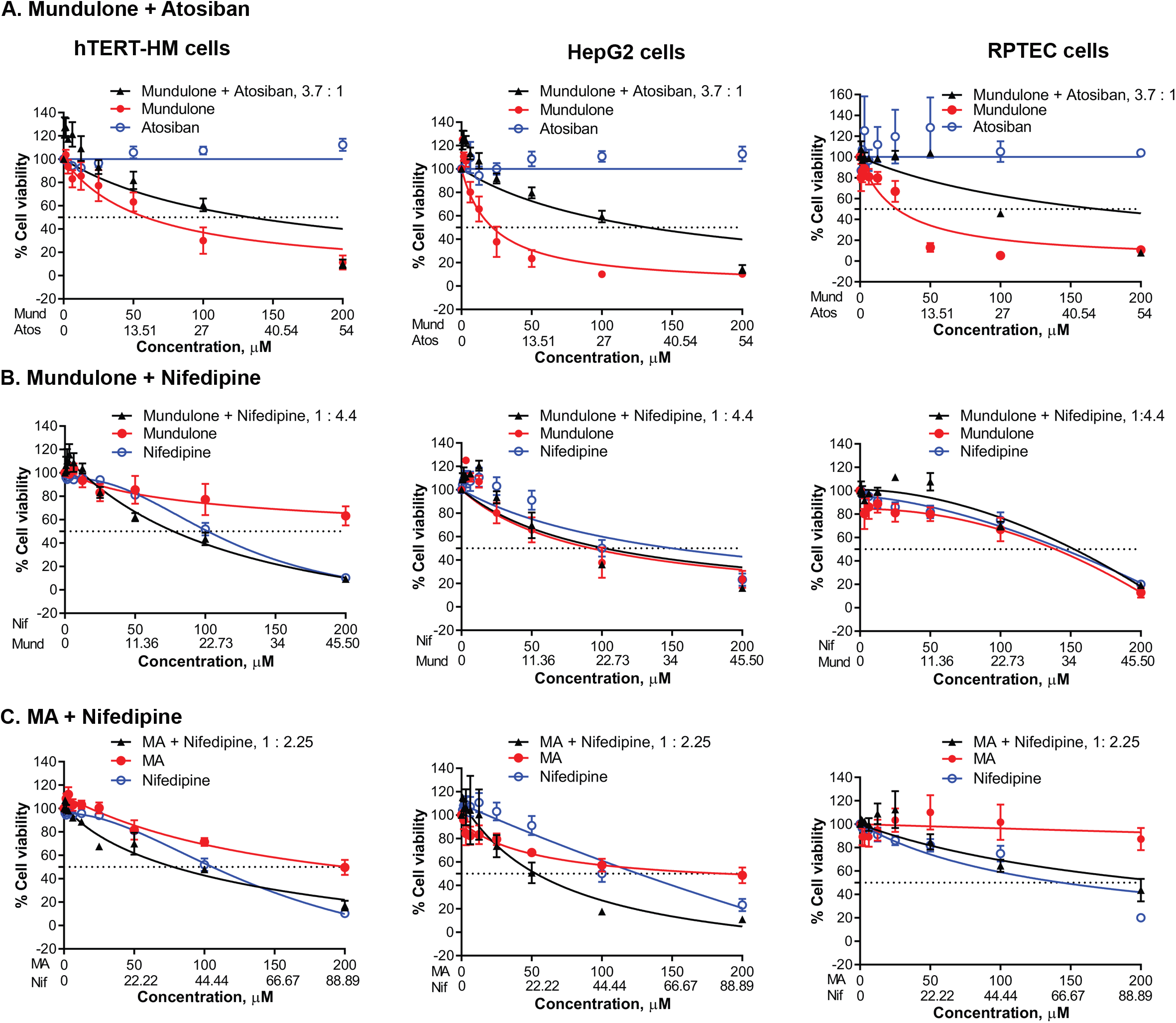
Cytotoxicity assessment of mundulone and MA synergistic combinations with clinical tocolytics. A WST-1 assay was used to examine the % cell viability of myometrial (hTERT-HM) cells, liver (HepG2) cells and kidney (RPTEC) cells after 72hr incubation with synergistic combinations of mundulone+atosiban (A), mundulone+nifedipine (B) and MA+nifedipine (C) at fixed ratios indicated on the graphs, and their respective single-compound controls. (Mund = Mundulone, Nif = Nifedipine, Atos = Atosiban, MA = Mundulone acetate. Non-linear regression was used to fit the data (Mean + SEM) and calculate IC_50_, which are provided in Table 3, along with p-values. A 2-way ANOVA with a post-hoc Tukey analysis was used to compare the E_max_ values (shown).

**Table 3:** Therapeutic index of mundulone and MA as single-agents as well as fixed-ratio combinations with current tocolytics.

| Drug combinations | hTERT-HM |  |  |  | HEPG2 |  |  |  | RPTEC |  |  |  | TI |
| --- | --- | --- | --- | --- | --- | --- | --- | --- | --- | --- | --- | --- | --- |
|  | Compound A<br>IC50 (μM) | Compound B<br>IC50 (mM) | Compound A;<br>Compound B |  | Compound A<br>IC50 (mM) | Compound B<br>IC50 (μM) | Compound A;<br>Compound B |  | Compound A<br>IC50 (μM) | Compound B<br>IC50 (μM) | Compound A;<br>Compound B |  |  |
|  |  |  | IC50 (mM) | p-value |  |  | IC50 (μM) | p-value |  |  | IC50 (μM) | p-value |  |
| Mundulone + Atosiban 3.7 : 1 | 58.9 | >200 | 133.9; 36.19 | 0.0038 | 22.1 | >200 | 132.4; 35.78 | <0.0001 | 26.02 | >200 | 169.2; 45.73 | <0.0001 | 10 |
| Mundulone + Nifedipine 1 : 4.4 | >200 | 106 | 19.15; 84.24 | <0.0001 | 92.6 | 150.2 | 23.16; 101.9 | 0.7152 | 119.5 | 143.9 | 40.25; 177.1 | 0.1978 | 2.81 |
| MA + Nifedipine 1 : 2.25 | 246.6 | 106 | 34.45; 77.52 | <0.0001 | 125.4 | 150.2 | 24.44; 55 | 0.0056 | >200 | 143.9 | 100; 225 | 0.0001 | 1.83 |

Compound A + Compound B values represent the concentration of compound A and compound B present in the combination at 50% inhibition of Ca<sup>2+</sup> mobilization or cell viability, respectively. Bolded values represent a favorable therapeutic index (TI = IC<sub>50</sub> Cell Viability Assay/IC<sub>50</sub> Ca<sup>2+</sup> Assay). p-values provided for drug combinations are for comparison between the combination and compound A.

### 3.5. *Ex vivo* tocolytic ability of synergistic combinations

Finally, we wanted to examine whether the synergistic combination, mundulone + atosiban with a favorable TI, exhibited synergistic tocolysis on term pregnant mouse myometrium in an *ex vivo* contractility assay. The concentration-dependent inhibition of tissue contractility by the fixed-ratio of mundulone + atosiban was compared to that of the individual compounds (Fig.6). The efficacy and potency of mundulone to inhibit mouse myometrial contractions significantly (p≤0.003) improved in combination with atosiban compared to use as a single agent (E_max_ = 69% and IC_50_ = 14uM versus E_max_ = 92% and 0.13uM, respectively).

**Figure 6.**
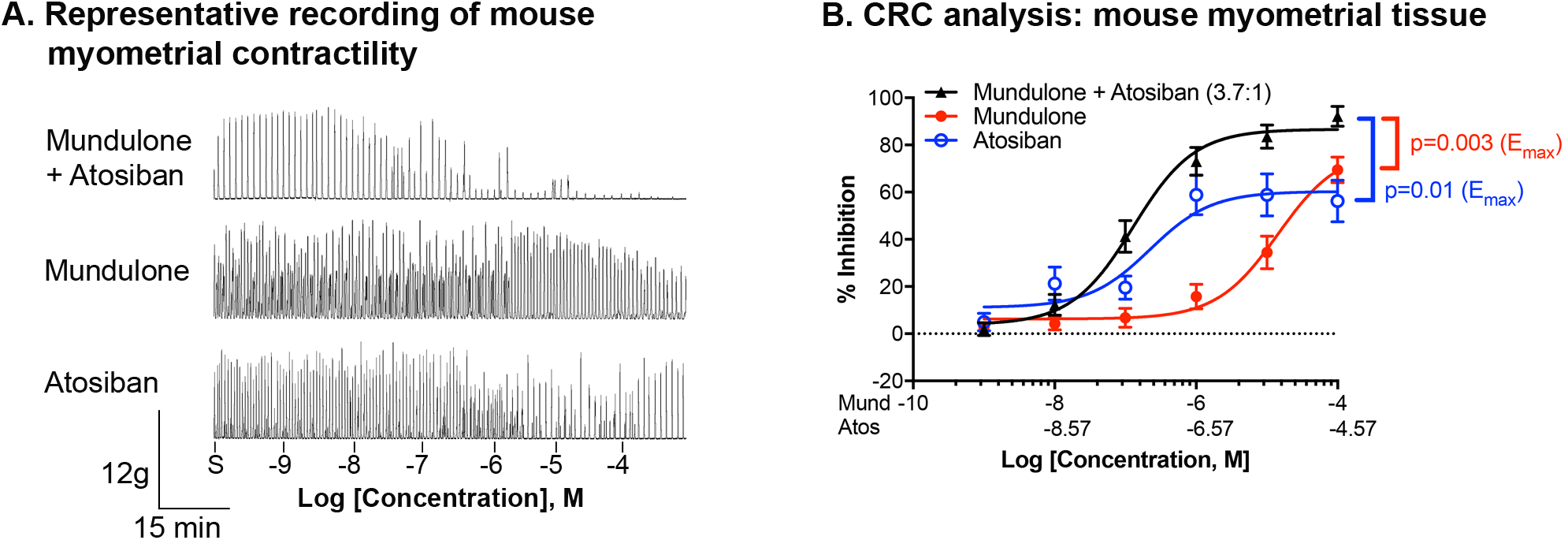
*Ex vivo* tocolytic effect of mundulone and atosiban combination. A. Representative recording of isometric spontaneous contractility (measured in grams of tension) of mouse myometrial tissue prior to the addition of increasing concentrations (10pm - 100uM) of mundulone +atosiban at a fixed ratio (3.7:1), as well as their single-compound controls (mundulone or atosiban). B. Recordings were analyzed for contractile AUC. Non-linear regression was used to fit the data (Mean ± SEM) and to calculate EC_50_ and E_max_. A 2-way ANOVA with a post-hoc Tukey analysis was used to compare the E_max_ values (shown).

## Discussion

It is important to continue efforts toward the discovery of novel tocolytics given the lack of clinically effective tocolytics without serious maternal and fetal side effects. Mundulone and its analog, MA, were previously identified in our high-throughput phenotypic screening assay of small-molecule compound libraries against intracellular-Ca^2+^ [3], given its function as the final common pathway controlling myometrial contractions [48, 49]. In the present study, we confirmed the ability of mundulone to inhibit *ex vivo* mouse and human uterine contractions in a concentration-dependent manner in an organ bath assay. Mundulone compared to MA displayed greater inhibitory efficacy on *in vitro* Ca^2+^-mobilization as well as *ex vivo* contractile activity. Other studies have similarly reported the uterine-relaxant effects of additional isoflavones and flavonoids: genistein, galetin 3,6-dimethyl ether, quercetin, and kaempferol [11, 12, 50].

Mundulone and other isoflavones have been reported to show hypotensive effects [51] and elicit vascular smooth muscle contractions, respectively. Due to this, as well as the previously stated off-target maternal cardiovascular side effects and fetal DA constriction of current off-label tocolytics, we tested the *in vitro* and *ex vivo* myometrial selectivity of mundulone and MA against aortic VSMCs and fetal DA, respectively. In our study, mundulone and MA exhibited *in vitro* selectivity to either potently or efficaciously inhibit Ca^2+^-mobilization from human primary myometrial cells compared to aortic VSMCs. The observed uterine-selectivity of mundulone was confirmed *ex vivo*, as this compound significantly inhibited myometrial contractility compared to vehicle control, without observed effects on fetal DA vasoreactivity. Thus, while mundulone displayed greater efficacy (1.8x), MA with an acetate group in position 3^//^ displayed greater myometrial selectivity in efficacy. Due to this, mundulone could benefit from structure activity relationship studies, as well as explored for use in combination with current clinically-utilized tocolytics. Moreover, expanded counter-screening efforts on other smooth muscle tissue types would expand our knowledge of the extent of mundulone’s and its analog’s uterine-selectivity.

Combination therapy is a promising approach to develop treatments for a wide range of disorders permitting lower doses that reduce off-target effects or increase therapeutic efficacy [52-57]. While combinations of current tocolytics (atosiban, nifedipine, indomethacin) with each other to identify synergy in potency and/or efficacy are reported, they are limited to examining a few fixed concentrations in small scale studies [58-62]. The current study is the first to report a high-throughput combination screen using a dose matrix approach that includes a wide range of concentrations for the discovery of novel tocolytic synergy. We identified *in vitro* synergistic combinations of isoflavone natural products with two current tocolytics. Mundulone exhibited synergism in potency and efficacy with atosiban, while showing only synergistic efficacy with nifedipine. MA had synergistic efficacy with only nifedipine. In our study, indomethacin was not found to have any synergistic effect with mundulone or MA.

When two drugs are combined, it is necessary to rule out unintended additive or synergistic toxicity early in the discovery process [63]. *In vitro* cell-based toxicity assays can be used to rapidly assess potential safety issues. *In vitro* therapeutic index obtained by measuring compound`s cytotoxicity is commonly used to quantify the extent of the safety at the desired efficacious conditions of a single- or combination of compounds. The toxicity of mundulone has been previously reported on HEK293 cells and zebrafish embryo[64, 65], however, mundulone acetate’s toxicity is not reported. In the current study, mundulone by itself was found to be toxic at low concentrations (TI = 0.8). However, when combined with atosiban the TI of mundulone dramatically improved to a favorable TI = 10, and therefore, shows potential for combination with current tocolytics. To this end, at least one other study reported improved inhibition of uterine contractility when another flavonoid containing plant extract from *Bryophyllum pinnatum* was combined with atosiban or nifedipine [18]. In our study, the *ex vivo* tocolytic efficacy and potency of mundulone in combination with atosiban significantly improved using term-pregnant mouse and human myometrial tissue.

Most published studies examining the tocolytic potential of natural products are of crude extracts, creating difficulty in pinpointing the activity to an isolated compound and limiting efforts for medicinal chemistry. The current study was conducted on a pure natural compound, providing an opportunity for further development. The mechanism pertaining to the tocolytic activity of mundulone and MA is not yet investigated. Mundulone and its structurally-related derivative compound, dihydromunduletone, are reported to inhibit the G protein coupled receptor GPR56’s activity [64]. Moreover, isoflavones are well known to be phytoestrogens with agonistic or antagonistic activity against estrogen receptors [66, 67]. Thus, it is worthwhile to probe the mechanism of both compounds, mundulone and MA, as it relates to modulation of intracellular Ca^2+^-regulated myometrial contractility through either a G protein coupled receptor, estrogen receptor and/or a different molecular target.

Collectively, this study demonstrates the tocolytic abilities of mundulone and MA through uterine-selective inhibition of intracellular-Ca^2+^. A novel *in vitro* high-throughput combination screen identified synergistic ratios of mundulone with atosiban with a favorable therapeutic index, whose tocolytic efficacy was validated in a separate *ex vivo* assay. Together, these data highlight that mundulone or its analogs warrant future development as single- and/or combination-tocolytic therapy for management of preterm labor.

## Acknowledgments

We would like to thank the nurses and physicians in the Department of Obstetrics and Gynecology at Vanderbilt University Medical Center for collecting myometrial biopsies, and all the women that kindly participated in this study. We thank the Vanderbilt Institute of Chemical Biology High Throughput Screening Facility for their technical assistance and use of their equipment for high-throughput assays. The Wave Front Biosciences Panoptic kinetic imaging plate reader was purchased with funds from the NIH Office of The Director S10OD021734. We also would like to thank Dr. Jennifer Condon (Wayne State University) for kindly providing the hTERT-HM cells used in this study.

## Figure Legends

**Supplementary Figure 1.**
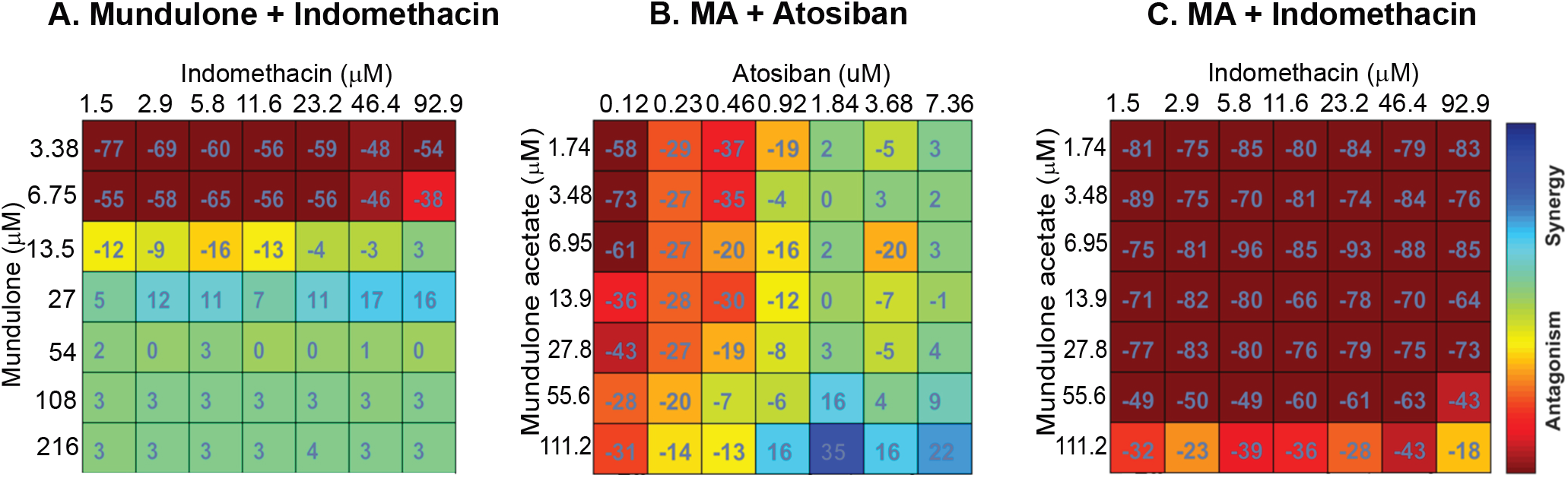
Combinations of mundulone and MA synergistic with clinical tocolytics that were not synergistic. A high-throughput Ca^2+^-assay using myometrial cells and mundulone or MA in combination with clinical tocolytics, indomethacin or atosiban, was performed. The %response data for each compound concentration was averaged and then analyzed with Combenefit software to provide synergy scores using the Bliss-independence model. Heat maps of synergy scores are shown for combinations of mundulone and MA with clinical tocolytics that did not display synergy: mundulone+indomethacin (A), MA+atosiban (B) and MA+indomethacin (C).

**Supplementary Figure 2.**
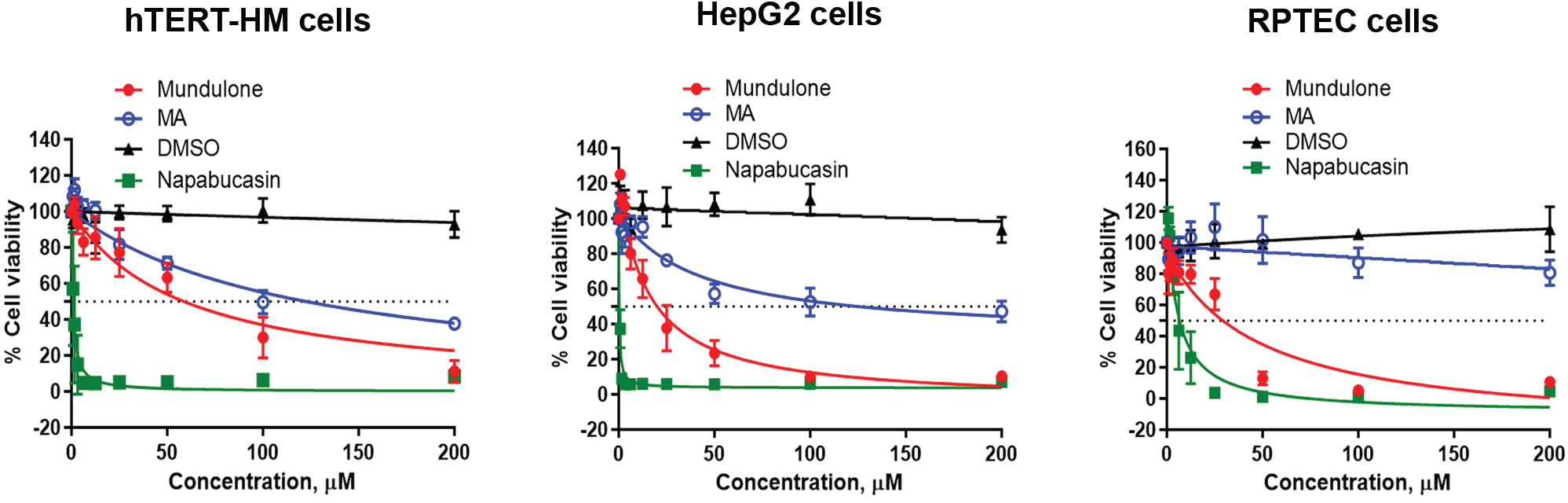
Effect of mundulone and MA on cell viability. A WST-1 assay was used to examine the % cell viability of myometrial (hTERT-HM) cells, liver (HepG2) cells and kidney (RPTEC) cells after 72hr incubation with either mundulone, MA, DMSO (vehicle control) or napabucasin (positive control, known-toxic compound). Non-linear regression was used to fit the data (Mean + SEM) and calculate IC_50_, which are provided in Suppl. Table 1, along with p-values and therapeutic indices.

---

**Supplemental Table 1:**
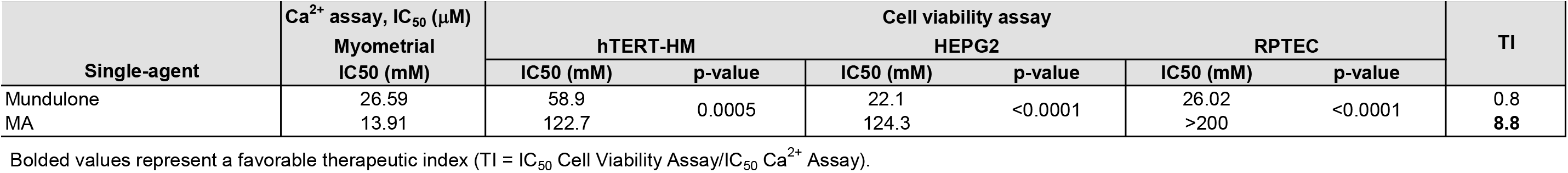
Therapeutic index of mundulone and MA as single-agents as well as fixed-ratio combinations with current tocolytics.

